# Molecular characterization of *Mycobacterium ulcerans* DNA gyrase and identification of mutations reduced susceptibility against quinolones *in vitro*

**DOI:** 10.1101/2021.09.29.462499

**Authors:** Hyun Kim, Shigtarou Mori, Tsuyoshi Kenri, Yasuhiko Suzuki

## Abstract

Buruli ulcer disease is a neglected necrotizing and disabling cutaneous tropical illness caused by *Mycobacterium ulcerans* (*Mul*). Fluoroquinolone (FQ), used in the treatment of this disease, has been known to act by inhibiting the enzymatic activities of DNA gyrase; however, the detailed molecular basis of these characteristics and the FQ resistance mechanisms in *Mul* remains unknown. This study investigated the detailed molecular mechanism of *Mul* DNA gyrase and the contribution of FQ resistance *in vitro* using recombinant proteins from the *Mul* subsp. shinshuense and Agy99 strains with reduced sensitivity to FQs. The IC_50_ of FQs against Ala91Vla and Asp95Gly mutants of *Mul* shinshuense and Agy99 GyrA subunits were 3.7- to 42.0-fold higher than those against wild-type enzyme. Similarly, the CC_25_ was 10- to 210-fold higher than those for the WT enzyme. Furthermore, the interaction between the amino acid residues of WT/mutant *Mul* DNA gyrase and FQ side chains was assessed via molecular docking studies. This is the first detailed study showing the contribution of *Mul* DNA GyrA subunit mutations to reduce the susceptibility against FQs.

## INTRODUCTION

Buruli ulcer disease (BU_D_) is an emerging chronic ulcerating illness caused by the environmental mycobacterium, *Mycobacterium ulcerans* (*Mul*), which primarily affects the skin, subcutaneous tissue, and occasionally bones. It is recognized by the World Health Organization (WHO) as a neglected tropical disease (1), and is the third most frequent skin mycobacterial disease worldwide after leprosy and tuberculosis (2, 3). BU_D_ is gradually increasing with approximately 2,000 to 5,000 new annual reported cases (4, 5). The cases have been reported in over 33 countries worldwide, primarily in tropical and subtropical regions (4), such as West Africa, Central Africa, and South Africa, and the Western Pacific countries (6-9). The reasons for increases in the past few years have not been understood (10-12).

The mode of transmission of *Mul* is not known although it is suggested to be through direct inoculation of the skin or subcutaneous tissue (13). *Mul* produces mycolactone, an immunomodulatory macrolide toxin which is the main pathogenic factor of BU_D_ (14). This toxin induces tissue necrosis, particularly in subcutaneous fat (14). Typically, *Mul* infections result in painless ulcers with undermined edges and necrotic sloughing which often affects the upper or lower limbs and the face (14). Recently, drug therapy against *Mul* has been administered through anti-mycobacterial antibiotics, including rifampicin-based combinations with either streptomycin, amikacin, or clarithromycin (15-17). Early and non-severe stages of BU_D_ can be treated with an 8-week regimen of rifampicin (10 mg/kg orally, once daily) combined with clarithromycin (7.5 mg/kg per body weight, twice daily), streptomycin (15 mg/kg intramuscularly, once daily), fluoroquinolone (FQ) or other antibiotics (15-19).

FQ is effective against *Mul in vitro* and *in vivo* (16, 20-22). Evidence exists that DNA topoisomerase II is the therapeutic target of the drug. The majority of eubacteria have two DNA topoisomerases II (DNA gyrase and DNA topoisomerase IV) which are essential for efficient DNA replication and transcription (23, 24), and among a few clinically validated targets for antibacterial therapies (25, 26). Remarkably, *Mul* expresses only DNA gyrase (27, 28) from a *gyrB* linked *gyrA* contig in the complete genome and this enzyme is the sole target of FQs (26). The catalytically active mycobacterial DNA gyrase has a GyrA_2_GyrB_2_ tetrameric structure (29) and is an ATP-dependent enzyme that transiently cleaves and unwinds double-stranded DNA to catalyze DNA negative supercoiling (30, 31). However, the detailed molecular mechanism of *Mul* DNA gyrase and the mechanisms of FQ resistance were not determined.

This study aimed to determine the functional analysis of *Mul* DNA gyrase activities *in vitro* from *Mul* shinshuense and Agy99 strains. DNA gyrase subunits of both strains were expressed and purified as a recombinant protein and its activity was investigated *in vitro* via supercoiling assays. In addition, specific structural interactions between wild-type (WT)/mutant *Mul* DNA gyrase and FQs were identified via molecular docking. Since the FQs tested had limited activity against the FQ-resistant *Mul* DNA gyrase, the development and design of novel antibiotics against BU_D_ are recommended.

## RESULTS

### Expression and purification of recombinant *Mul* DNA gyrases

The entire gene sequences of WT *gyrA* and *gyrB* from *Mul* shinshuense and Agy99 strains and mutant *gyrA* (Ala91Val and Asp95Gly) were amplified and inserted into expression vector pCold-I on downstream of the *cspA* promoter to heterologous express N-terminal hexahistidine-tagged gyrase subunits. Molecular docking predicted that the his_6_-tag was located away from the FQ binding site of the *Mul* DNA GyrA (Fig. 1A) and GyrB (data not shown) subunit suggesting that it will not interfere with GyrA activity. Expressed WT/mutant GyrA and GyrB subunits were purified to homogeneity using a two-step column chromatographic procedure described in the Materials and Methods section with the expected molecular masses of GyrA (93 kDa) and GyrB (76 kDa) subunits determined by sodium dodecyl sulfate-polyacrylamide gel electrophoresis (SDS-PAGE) (Fig. 1B). All recombinant DNA gyrase subunits were obtained at high purity (> 95%) in milligram amounts. Contaminating *Escherichia coli* (*E. coli*) topoisomerase activity was denied by the lack of supercoiling activities either only with *Mul* GyrA or GyrB subunit (Fig. 2, lanes 2 and 3 for *Mul* shinshuense, and lanes 6 and 7 for *Mul* Agy99).

**Fig. 1.**
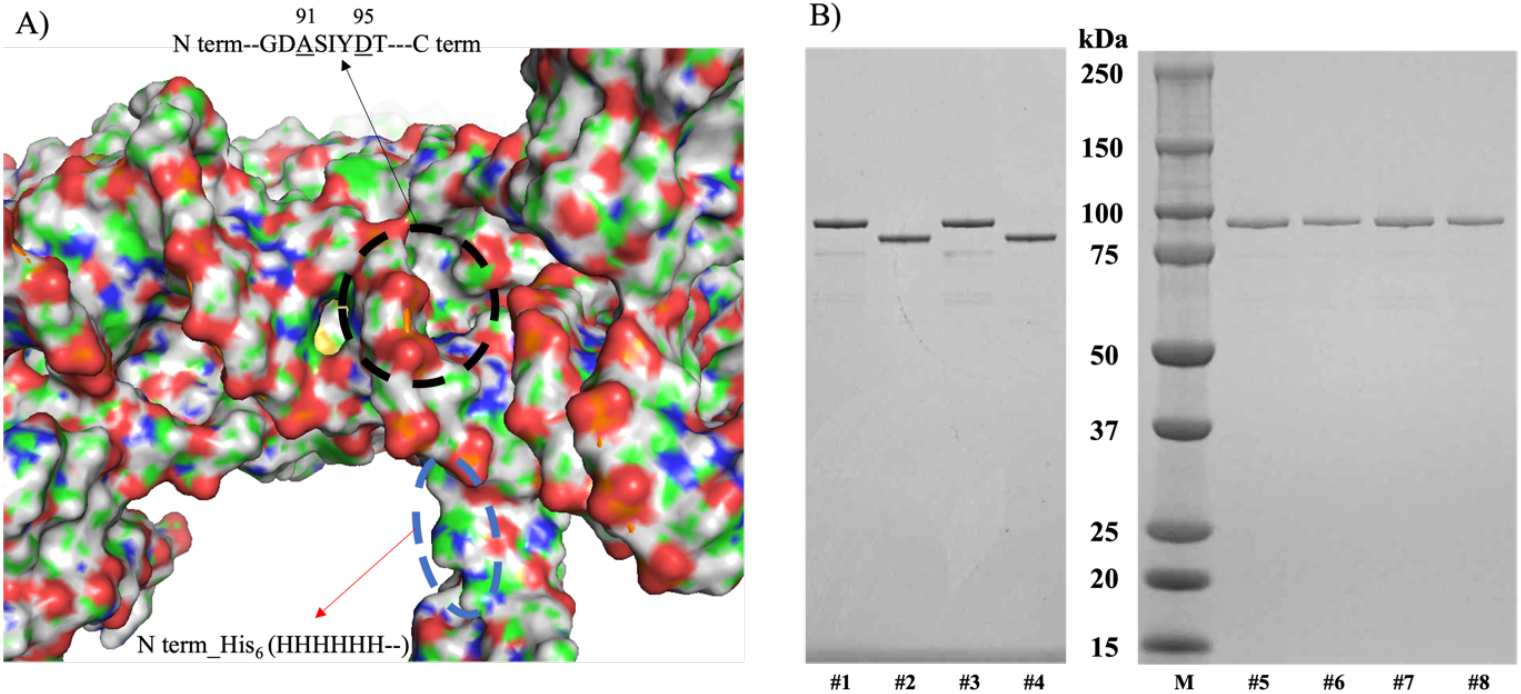
Purity of recombinant *Mul* DNA gyrase subunits. A) Black and blue circles represent the FQ binding domain and the location of the N-terminus his_6_-tag, respectively. B) 5-20% SDS-PAGE (ATTO, Tokyo, Japan), of WT and mutant *Mul* DNA gyrase subunits from shinshuense and Agy99 strains. Approximately 3 μM of each subunit was loaded into each well. Lanes 1 and 2: WT *Mul* shinshuense GyrA and GyrB subunit, respectively; lanes 3 and 4: WT *Mul* Agy99 GyrA and GyrB subunits, respectively. Lane M, size markers (kDa, Bio-Rad Lab. Inc., Japan); lanes 5 and 6: Ala91Val and Asp95Gly *Mul* shinshuense GyrA mutants, respectively; lanes 7 and 8: Ala91Val and Asp95Gly *Mul* Agy99 GyrA mutants, respectively.

**Fig. 2.**
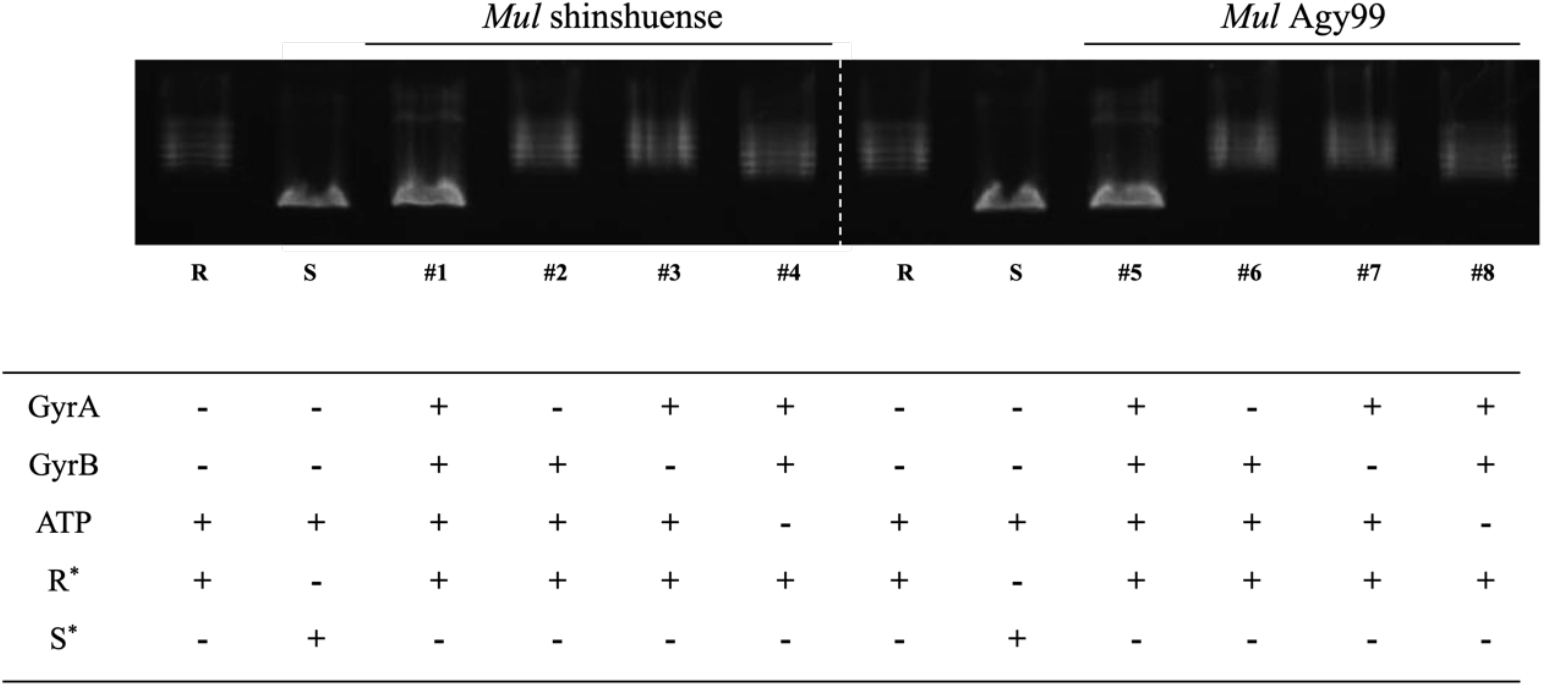
ATP-dependent DNA supercoiling activities of WT *Mul* DNA gyrases. Three μM GyrA and 1 μM GyrB (*Mul* shinshuense), and 3 μM of GyrA and GyrB (*Mul* Agy99) in the presence or absence of ATP. Lanes 1 and 5: GyrA and GyrB from shinshuense and Agy99 strains, respectively; lanes 2 and 6: absence of GyrA subunit; lanes 3 and 7: absence of GyrB subunit; lane 6: absence of ATP. R^*^ and S^*^ denote relaxed and supercoiled pBR322 DNA, respectively.

### Supercoiling activities of WT and mutant *Mul* DNA gyrase

The supercoiling activities of each DNA gyrase subunit were investigated by varying the subunit concentration (Fig. 3). Three µM GyrA and 1 µM GyrB (for WT *Mul* shinshuense, Fig. 3A), or 3 µM of GyrA and GyrB (for mutants shinshuense and WT and mutants Agy99, Fig. 3B-D) converted 100% of 0.3 µg of relaxed pBR322 plasmid DNA substrate to its supercoiled form. This subunit concentration was used for all subsequent enzyme assays.

**Fig. 3.**
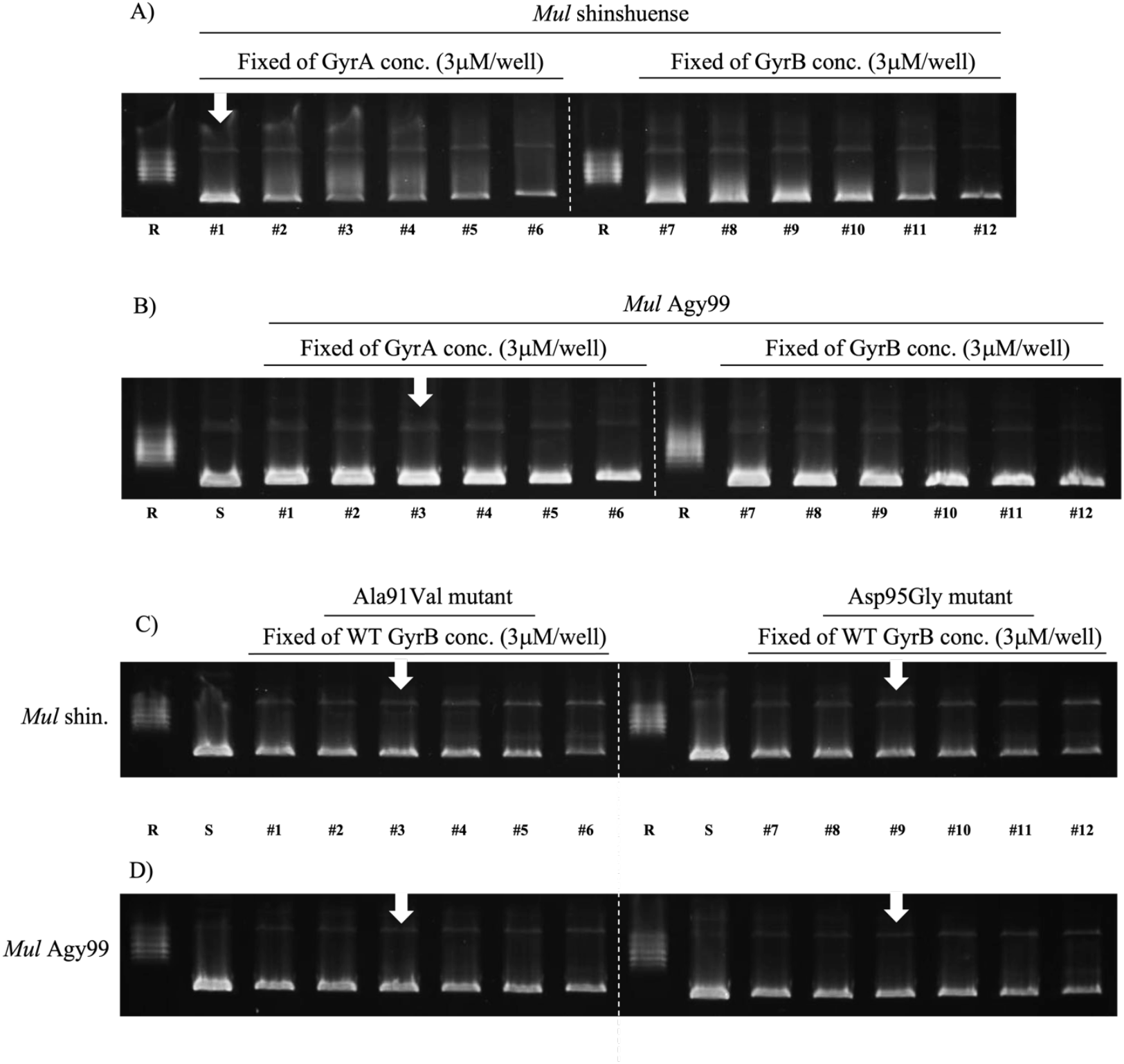
Concentration-dependent DNA gyrase supercoiling assays. Assays were performed using a fixed concentration (3 μM) of GyrA (left panel) with various concentrations of GyrB (1, 2, 3, 6, 12, and 24 μM) or fixed GyrB with variable GyrA concentrations (right panel). A) WT *Mul* shinshuense; B) WT Agy99; C) Ala91Val and Asp95Gly *Mul* shinshuense mutants; D) Ala91Val and Asp95Gly *Mul* Agy99 mutants. #1 and 7, 1 μM; #2 and 8, 2 μM; #3 and 9, 3 μM; #4 and 10, 6 μM; #5 and 11, 12 μM; #6 and 12, 24 μM, respectively. The optimum levels of DNA gyrase subunits are denoted by white arrows.

The supercoiling activity of *Mul* DNA gyrases was ATP-dependent (Fig. 2, lane 4 and 8) and required the combination of GyrA and GyrB subunits (Fig. 2, lane 1 and 5). Mutant GyrA (Ala91Val and Asp95Gly) had DNA supercoiling activity in the presence of WT GyrB (data not shown). Furthermore, we found that the optimum temperature of *Mul* DNA gyrase was 30–37 °C and its activity decreased at 40 °C (Fig. 4) and is similar to those of the *Mycobacterium leprae* DNA gyrase from our previously reported (34); therefore, all other assays were performed at 30 °C.

**Fig. 4.**
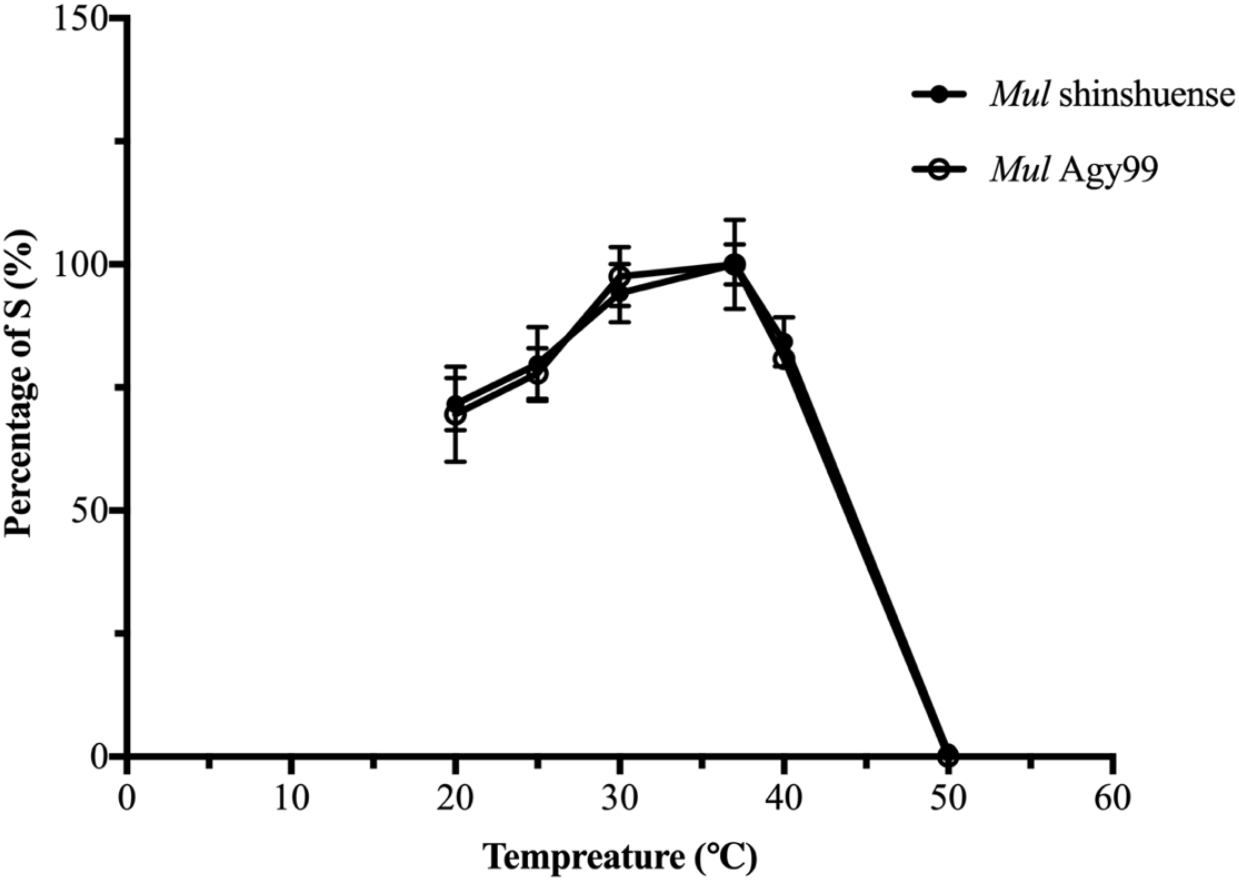
Temperature-dependent *Mul* shinshuense and Agy99 DNA gyrases supercoiling activities. Assays were performed at 20 °C, 25 °C, 30 °C, 37 °C, 42 °C, and 50 °C using 3 μM GyrA and 1 μM of GyrB for *Mul* shinshuense (•), and 3 μM *Mul* Agy99 GyrA and GyrB (○). Electrophoresis results are shown below the graph. Assays were performed in triplicate. S represents supercoiled pBR322 DNA.

### Inhibitory effect of FQs against *Mul* WT and mutant DNA gyrases

The inhibitory effect of FQs ciprofloxacin (CIP), moxifloxacin (MOX), and levofloxacin (LVX) on each WT and mutant (*Mul* shinshuense and Agy99) DNA gyrase was elucidated using the DNA supercoiling assay (Fig. 5). IC_50_ values were ordered from low to high (Table 2), with the structure of each FQ shown in Table 2A-D. IC_50_s from both strains of DNA gyrase were comparable to that observed (Table 2). The mutant DNA gyrase was highly resistant to inhibition by CIP and LVX (Fig. 5 and Table 2) with IC_50_s of >320 µg/mL, whereas the *Mul* shinshuense and Agy99 WT gyrase were 11.80 and 7.52 µg/mL, respectively (Table 2). To examine the effects of FQ on cleavage complex formation by *Mul* recombinant DNA gyrases, cleavage activities were performed in which supercoiled pBR322 DNA was incubated with WT or mutant DNA gyrases in the presence or absence of increasing concentrations of FQs. The representative results of cleavage activity using LVX against *Mul* Agy99 DNA gyrase were shown in Fig. 6., and Table 2 presents the CC_25_ of CIP and MOX. The CC_25_ of FQs for WT DNA gyrase ranged from 0.038 to 1.53 μg/mL, while those for the mutant DNA gyrases ranged from 1.04 to 67.68 μg/mL (Table 2).

**Fig. 5.**
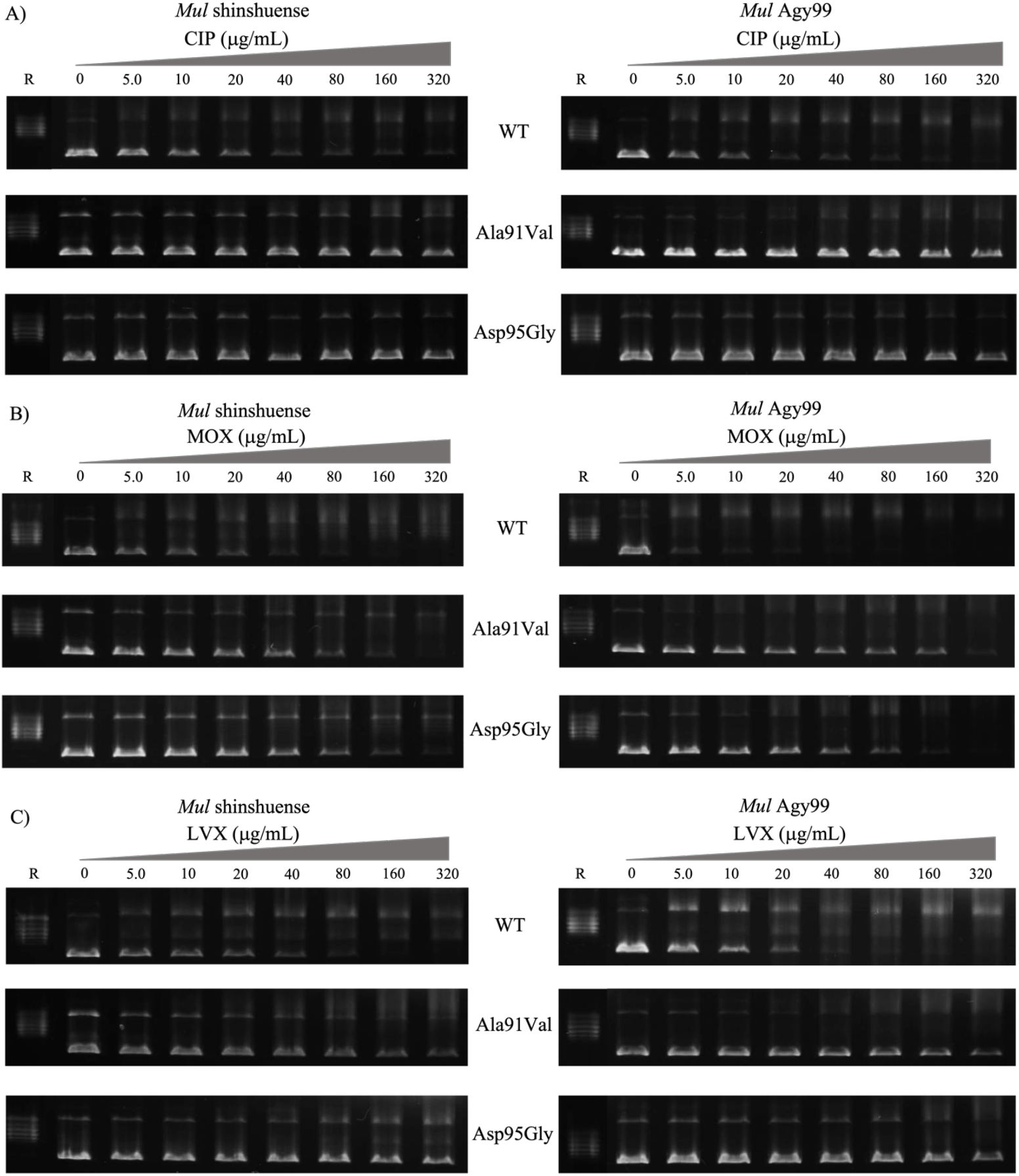
Inhibition of WT and mutant *Mul* DNA gyrase supercoiling activities by FQs. Optimum concentrations of DNA gyrase subunits in the presence or absence of the indicated amounts (μg/mL) of (A) CIP, (B) MOX, and (C) LVX. Inhibition of WT/mutant *Mul* shinshuense and Agy99 DNA gyrase was shown in the left and right panels, respectively. R denotes relaxed pBR322 DNA.

**Table 1.**
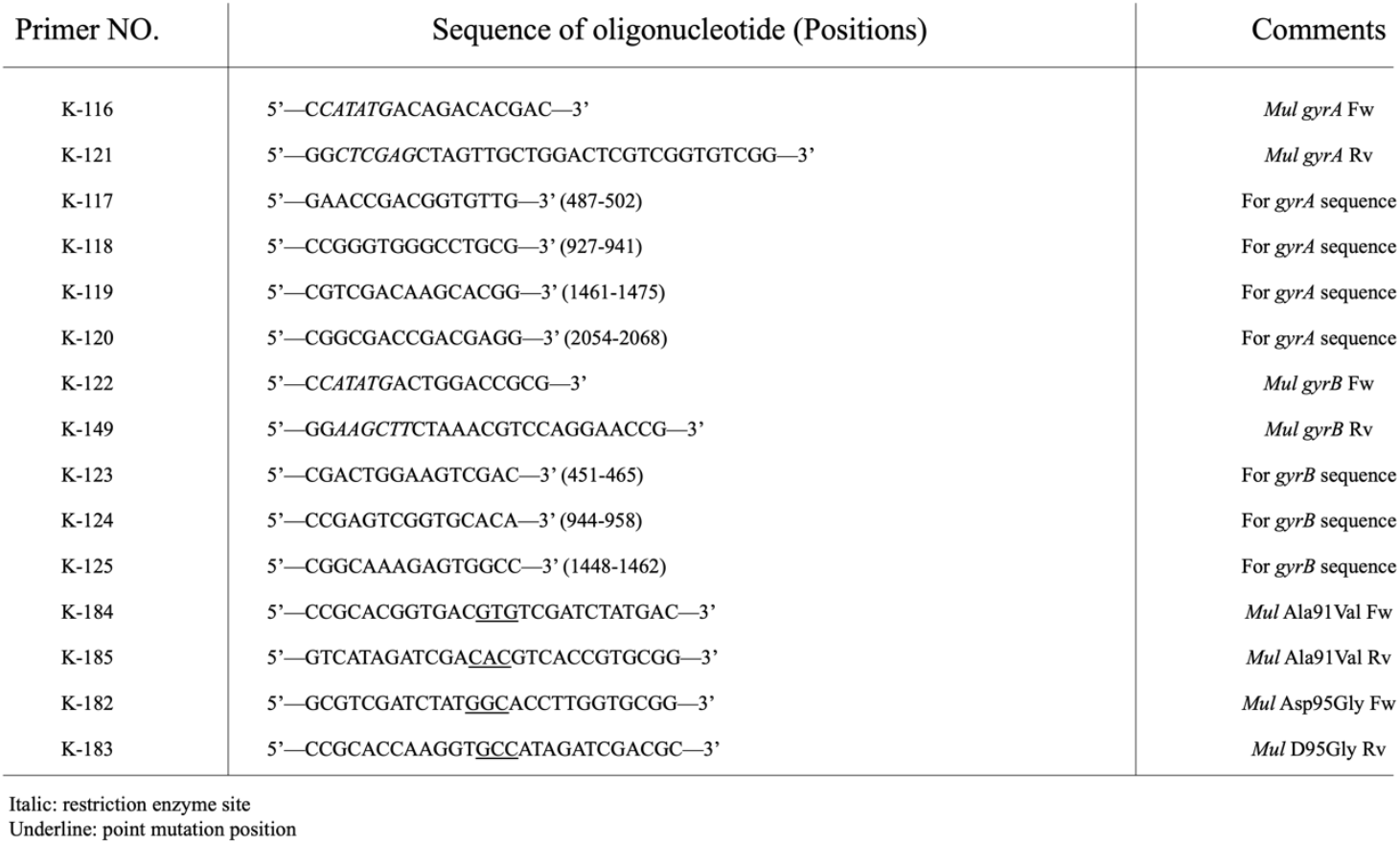
List of oligonucleotides.

**Table 2.**
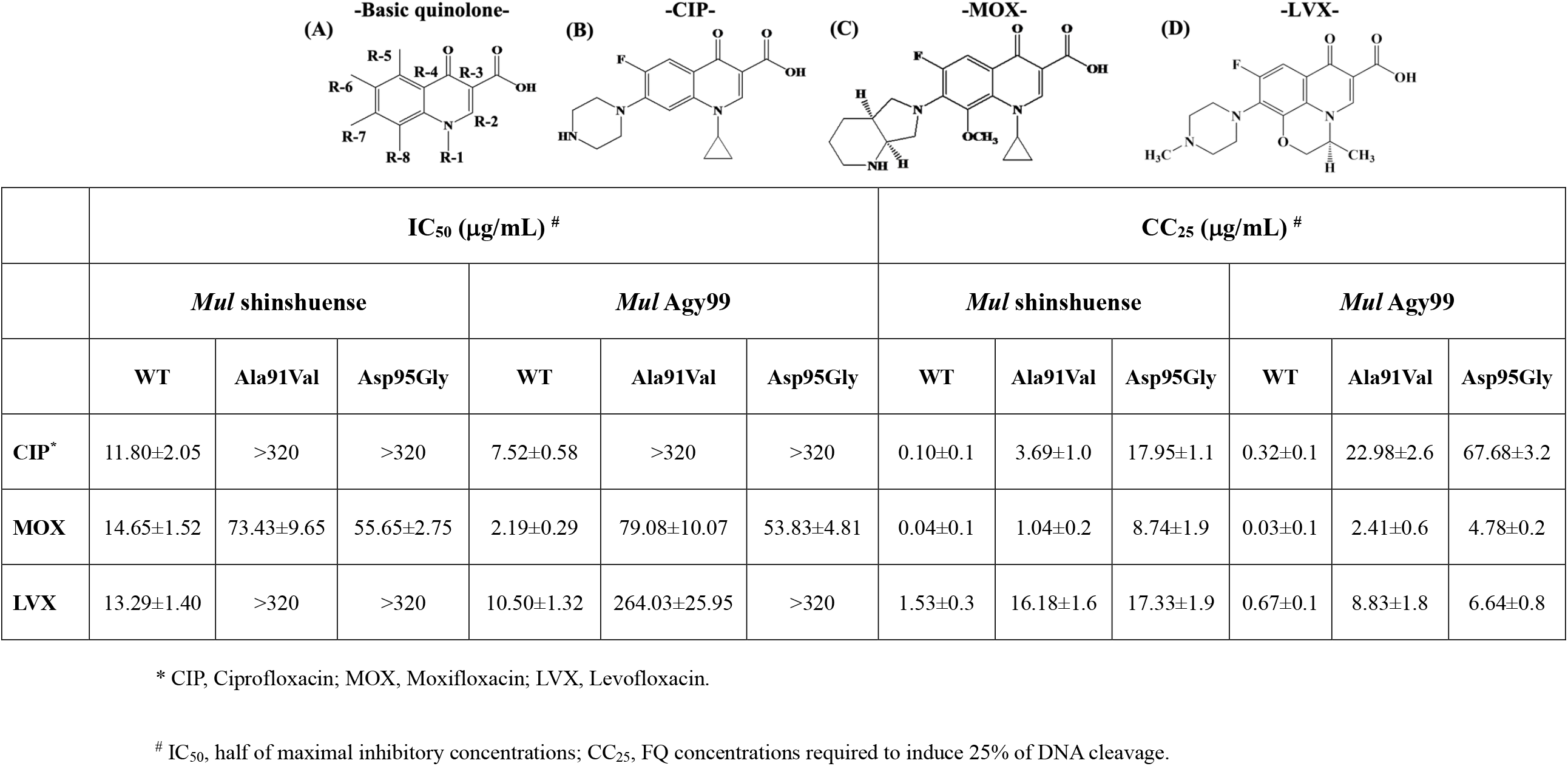
The results of inhibition assay and structure of FQs.

|  | IC<sub>50</sub> (μg/mL) # |  |  |  |  |  | CC<sub>25</sub> (μg/mL) # |  |  |  |  |  |
| --- | --- | --- | --- | --- | --- | --- | --- | --- | --- | --- | --- | --- |
|  | *Mul shinshuense* |  |  | *Mul Agy99* |  |  | *Mul shinshuense* |  |  | *Mul Agy99* |  |  |
|  | WT | Ala91Val | Asp95Gly | WT | Ala91Val | Asp95Gly | WT | Ala91Val | Asp95Gly | WT | Ala91Val | Asp95Gly |
| **CIP\*** | 11.80±2.05 | >320 | >320 | 7.52±0.58 | >320 | >320 | 0.10±0.1 | 3.69±1.0 | 17.95±1.1 | 0.32±0.1 | 22.98±2.6 | 67.68±3.2 |
| **MOX** | 14.65±1.52 | 73.43±9.65 | 55.65±2.75 | 2.19±0.29 | 79.08±10.07 | 53.83±4.81 | 0.04±0.1 | 1.04±0.2 | 8.74±1.9 | 0.03±0.1 | 2.41±0.6 | 4.78±0.2 |
| **LVX** | 13.29±1.40 | >320 | >320 | 10.50±1.32 | 264.03±25.95 | >320 | 1.53±0.3 | 16.18±1.6 | 17.33±1.9 | 0.67±0.1 | 8.83±1.8 | 6.64±0.8 |

\* CIP, Ciprofloxacin; MOX, Moxifloxacin; LVX, Levofloxacin.

**Fig. 6.**
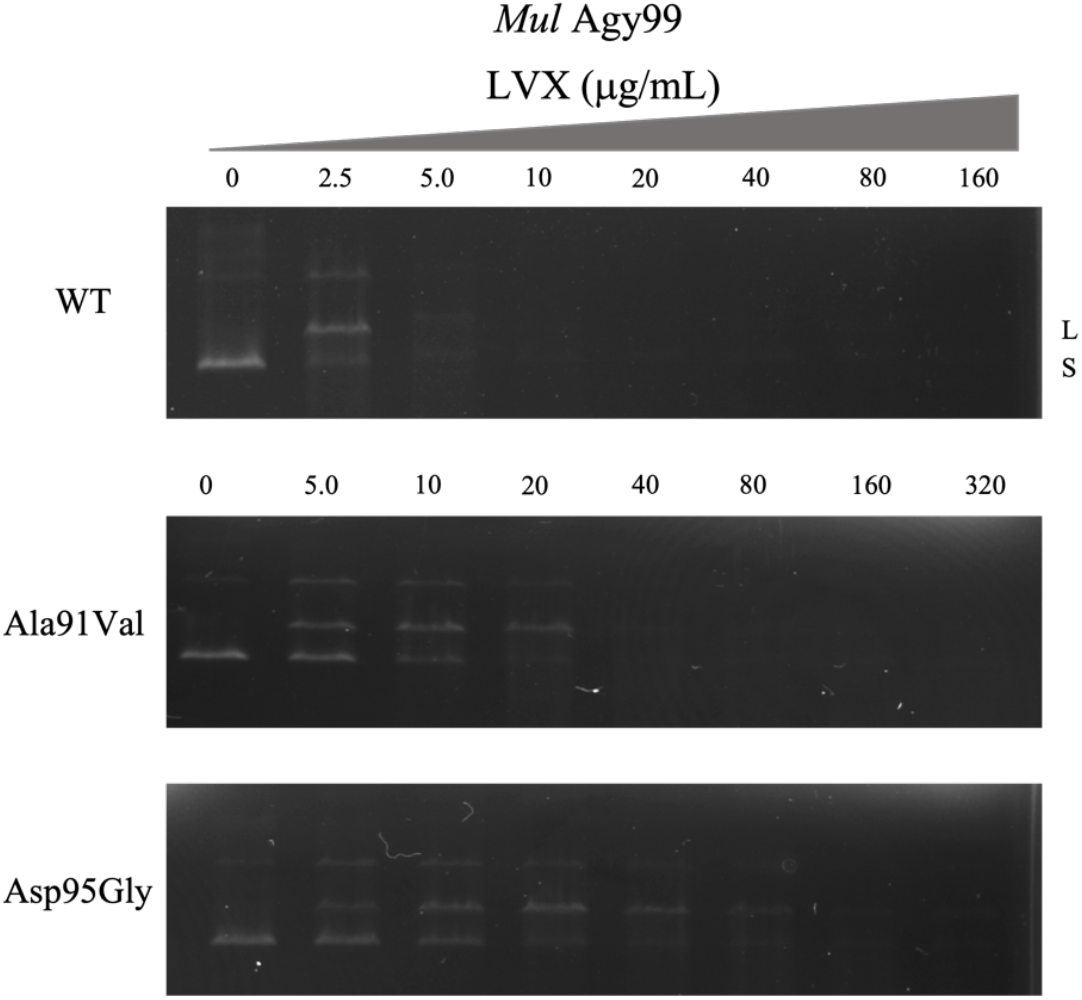
CIP-mediated DNA cleavage complex by WT and mutants *Mul* shinshuense DNA gyrases. Supercoiled pBR322 DNA (0.1 μg) was incubated with each WT and mutants *Mul* DNA gyrases in the presence of increasing LVX concentrations indicated (0–320 μg/mL). After the addition of each 3 μL of 2% SDS and 1 mg/mL proteinase K for 30 min at 37 °C, and then the reactions were stopped, and the mixture samples were analyzed by electrophoresis in 1% agarose gels. S and L denote supercoiled and linearized pBR322 DNA, respectively.

### Binding mode between DNA gyrase and FQs

The detailed interaction between WT/mutant *Mul* shinshuense DNA gyrase and FQs was determined via molecular docking using molecular operating environment (MOE) software (Fig. 7). The representative molecular docking results of CIP or MOX against *Mul* WT shinshuense and mutants DNA gyrase are shown in Fig. 7, whereas the Agy99 DNA gyrase data are not shown because the amino acid sequence has 95% homologous identity and FQ binding site was 100%. The docking score and root mean square deviation (RMSD) between CIP and WT DNA gyrase, or GyrA mutants (Ala91Val/Asp95Gly) were −6.3081 and 4.02 Å, or −6.3239/−6.4195 and 3.48/4.04 Å, respectively (Fig. 7C); Meanwhile, the docking score and RMSD between MOX and WT DNA gyrase, or GyrA mutants (Ala91Val/Asp95Gly) were −8.0615 and 2.15 Å, or −7.9429/−7.6337 and 1.80/3.35 Å, respectively (Fig. 7C). A hydrogen bonding network between the side chains of CIP and the amino acids of WT GyrA subunit: Ala91 (2.53 and 2.73 Å), Asp95 (2.94 and 2.82 Å), and Thr96 (2.44 Å) was observed (Fig. 7A). However, the interaction of CIP with the Asp95Gly GyrA mutant was not observed (Fig. 7A) which correlated with the significantly higher IC_50_ value of the Asp95Gly GyrA mutant compared with that of the WT and Ala91Val mutants. Similarly, the binding mode of MOX and distance were observed (Fig. 7B). Based on the molecular docking results, these amino acid residues in the GyrA subunit may play an important role in the interaction between *Mul* DNA gyrase and FQs.

**Fig. 7.**
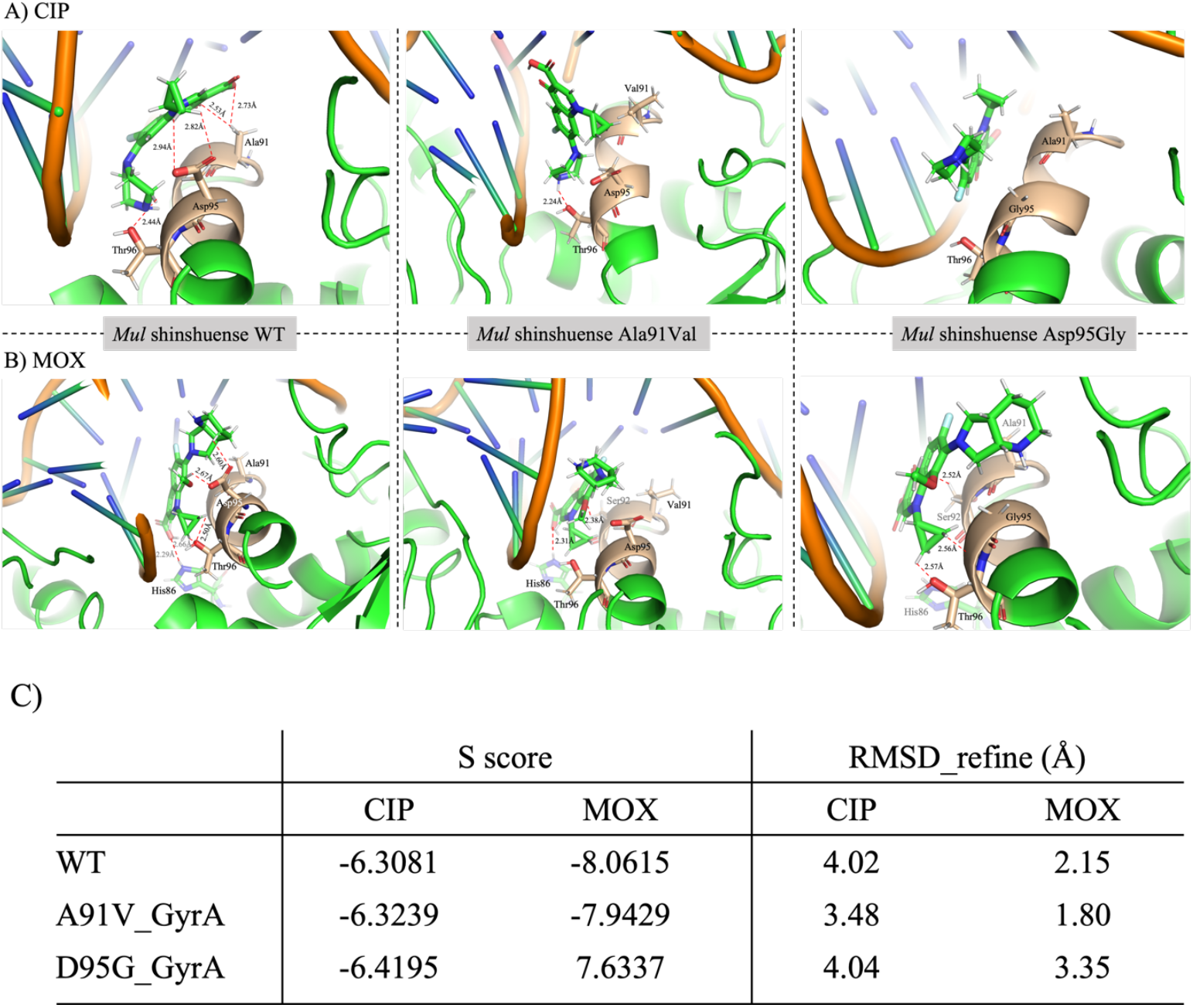
CIP binding mode in WT and mutant *Mul* shinshuense DNA gyrase A subunit. Molecular docking studies of WT and mutant *Mul* DNA gyrase with FQs were performed using the MOE software. The binding modes of CIP (A) and MOX (B) were shown within the catalytic site of *Mul* shinshuense DNA GyrA and the mutated amino acids (Ala91Val and Asp95Gly). Amino acids 90 to 96 were depicted in wheat color. The dotted red line indicates hydrogen bonding, and the distance between the amino acid residues and a side chain of FQs is indicated.

## DISCUSSION

Although the inhibitory effects of FQs against BU_D_ are known *in vitro* and *in vivo* (16, 20-22), the molecular details between *Mul* DNA gyrase and FQs interactions are not understood. Nakanaga K. *et al*. (32) reported the drug susceptibility test of *Mul* shinshuense and Agy99 strain, which results were showed the different MIC data between *Mul* shinshuense (0.25 μg/mL) and Agy99 (8.0 μg/mL) strains for LVX. In this regard, we are focused on the more detailed mechanism of FQ resistance against two strains *in vitro* using a bacterial recombinant system. Furthermore, we investigated the amino acid substitutions at positions 91 and 95 on the GyrA subunit from both *Mul* strains which is equivalent to positions 90 and 94 in *M. tuberculosis* known to contribute to FQ resistance (33-35). The resistance of mutant DNA gyrase to FQs was demonstrated at the molecular level using purified recombinantly expressed subunits by supercoiling and inhibition assays.

To measurement of absolute concentration for DNA gyrase activity, the concentration of purified DNA gyrase subunits was calculated by two steps, because the DNA gyrase activity was required the 1:1 ratio of GyrA and GyrB subunit. The first step was a general calculator system from the Qubit assay kit (data not shown), and the second step was a concentration-dependent supercoiling assay with variable concentrations of GyrA or GyrB subunits (Fig. 3). The optimum concentrations of GyrA and GyrB subunits (1:1 ratio) were decided; WT *Mul* shinshuense GyrA and GyrB subunits were 3 μM and 1 μM (the first step value indicated), respectively., and *Mul* Agy99 and mutants were each 3 μM of GyrA and GyrB subunit (Fig. 3).

The amino acid substitution of GyrA subunits at Ala91Val and Asp95Gly showed reduced sensitivity to all FQs, and more markedly than for WT DNA gyrase (2.19–14.65 μg/mL) (Fig. 5, 6 and Table 2). MOX exhibited the lowest FQ sensitivity against *Mul* GyrA mutants. The IC_50_ values of CIP, MOX, and LVX were over 20-fold higher against *Mul* shinshuense DNA gyrase mutants than those for WT DNA gyrase (Fig. 5 and Table 2). Similar results were observed for *Mul* Agy99 GyrA mutants (Fig. 5 and Table 2) and this may be related to its amino acid sequence homology (95%) to the *Mul* shinshuense GyrA subunit (data not shown). Furthermore, to examine the effects of quinolone on cleavage complex formation by *Mul* recombinant DNA gyrases were performed that a similar tendency was observed in the DNA cleavage activities, CC_25_ was over 20-fold higher than for the WT DNA gyrase (Fig. 6 and Table 2). The observations suggested the contribution of amino acid substitutions Ala91Val and Asp95Gly in the GyrA subunit resulted in reduced sensitivity to FQs and led to the emergence of *Mul*.

The structure-activity relationship between WT/mutant DNA gyrases and FQs was analyzed by molecular docking using MOE software (Fig. 7) to predict whether the GyrA subunit is associated with FQ resistance. A prediction of the three-dimensional (3D) structure of *Mul* shinshuense GyrA subunit was generated for structure-binding analysis with FQs because the crystal structure has not been reported. The WT or mutant *Mul* shinshuense GyrA subunit was modeled by Swiss-Model (https://swissmodel.expasy.org) using PDB 3IFZ (36) and 6RKS (37) as templates (Fig. 7). The position of amino acid residues Ala91, Asp95, and Thr96 in WT GyrA interacted with that of the R3, R1, and R7 ring of CIP, respectively (Fig. 7A left panel), which is suggested to contribute to potent inhibitory activity against *Mul* WT DNA gyrase (IC_50_ of 11.80 µg/mL). Modeling showed that CIP tightly binds to the quinolone-binding site on the GyrA subunit through hydrogen bonding interactions (CH_3_---COO [R3], COO---CH_2_ [R1], and OH---CH_2_ [R7] (Fig. 7A left panel and Table 2). In contrast, the efficacy of CIP against the Asp95Gly mutant on the GyrA subunit was significantly lower (IC_50_ >320 µg/mL) than that of the WT and Ala91Val mutant (Fig. 7A right panel and Table 2). The loss of hydrogen bonding may induce substantial conformational changes in the GyrA subunit which disrupts enzymatic activity. Modeling of the MOX ligand in place of CIP showed similar interactions (Fig. 7B). In summary, the two mutated amino acids (Ala91Val or Asp95Gly) likely play important roles in causing reduced sensitivity to FQ through altering the hydrogen bonding network in *Mul* DNA gyrase.

In conclusion of in this study, Ala90Val and Asp95Gly amino acid substitutions in *Mul* DNA gyrase reduced sensitivity to FQs *in vitro*. The present findings will aid in the design and development of novel BU_D_ antibiotics against the possible future emergence of FQ-resistant *Mul* carrying GyrA with these amino acid substitutions.

## MATERIALS AND METHODS

### Materials

CIP, LVX (LKT Laboratory, Inc., Minnesota, USA), and MOX (Toronto Research Chemicals Inc.) were used for FQ inhibition assays, while ampicillin (Wako Pure Chemicals Ltd., Tokyo, Japan) was used for culturing *E. coli* harboring plasmids. Oligonucleotides were synthesized by Eurofins Genomics Inc. (Tokyo, Japan). A TOPO TA cloning kit (PCR^®^ 4-TOPO^®^) was purchased from Life Technologies (Carlsbad, CA, USA) and used for cloning and nucleotide sequencing. DNA electrophoresis chemicals and restriction endonucleases were obtained from New England BioLabs, Inc. (Ipswich, MA, USA). Relaxed pBR322 DNA was purchased from John Innes Enterprises Ltd. (Norwich, United Kingdom). Protease inhibitor cocktail (Complete Mini, EDTA-free) was purchased from Roche Applied Science (Mannheim, Germany).

### Bacterial strains and plasmids

Genomic DNA from the *Mul* shinshuense (28) and Agy99 strains (27) was gifted from Dr. Yuji Miyamoto, Leprosy Research Center, National Institute of Infectious Diseases (Tokyo, Japan). *E. coli* Top 10 and DH5α strains were used as hosts for molecular cloning. The pCold-I vector (Takara Bio Inc., Shiga, Japan) was used to construct an expression vector to produce WT and mutant versions of recombinant GyrA and GyrB proteins from *Mul* shinshuense and Agy99 strains. *E. coli* BL21 (DE3) (Merck KGaA, Darmstadt, Germany) was used for protein expression.

### Construction of WT *gyrA* and *B* expression plasmids

WT *gyrA* and *gyrB* genes were amplified from genomic DNA of *Mul* shinshuense and Agy99 strain via PCR. The reaction mixtures (20 µL) contained 10× LA PCR buffer II (Mg^2+^-free), 2.5 mM dNTP mixture, 2.5 mM MgCl_2_, 250 ng genomic DNA from *Mul* shinshuense or Agy99 strains, 1.25 units of LA Taq polymerase (Takara Bio Inc.), and 0.1 μM of each primer. The primer details are listed in Table 1. PCR was conducted using a Takara PCR thermal cycler Dice mini (Takara Bio Inc.) as follows: denaturation at 98 °C for 2 min, 35 cycles of denaturation at 98 °C for 10 s, annealing at 60 °C for 10 s, and extension at 72 °C for 3 min 30 s, with a final extension at 72 °C for 2 min. The PCR products corresponding to the 2.5-kb *gyrA* and 2.1-kb *gyrB* fragments were ligated into the TA cloning plasmid, transformed into *E. coli* Top 10 and plated onto Luria–Bertani (LB) agar containing ampicillin (100 µg/mL). Colonies were selected, and plasmids were purified using a Miniprep DNA purification kit (Promega Madison, WI, USA), followed by digestion with *Nde*I and *Xho*I (for *gyrA*)/ *Hind*III (for *gyrB*). The *gyrA* and *gyrB* fragments were ligated into the pCold-I expression vector restriction digested with the same restriction endonucleases. Mutant *Mul* DNA gyrases (Ala91Val or Asp95Gly) were generated from WT *gyrA* and *gyrB* using a QuickChange site-directed mutagenesis kit (Agilent Technologies, Inc., Santa Clara, CA, USA) according to the manufacturer’s instructions using the primers stated in Table 1. Plasmids were purified using a Miniprep DNA purification kit. WT and mutant plasmids were confirmed by sequencing (GENEWIZ corp., Tokyo, Japan) and were checked for errors by comparing to their respective WT and mutant sequences using BioEdit software 7.0.5.3 (http://www.bioedit.com/).

### Overexpression and purification of recombinant *Mul* DNA gyrase subunits

Recombinant DNA gyrase subunits were purified as previously described (38-40) with minor modifications. Briefly, the recombinant WT and mutant *Mul gyrA* and *gyrB* expression vectors were transformed into *E. coli* BL21 (DE3) cells. Single colonies were picked and grown overnight at 37 °C in 4 mL of LB medium containing 100 µg/mL ampicillin. Overnight cultures were used to inoculate 400 mL of fresh LB medium with ampicillin. Cells were cultured at 37 °C for 7-8 h until the optical density (OD) at 600 nm reached 0.6 to 0.8, followed by the addition of 1 mM isopropyl β-D-1-thiogalactopyranoside (Wako Pure Chemicals Ltd.) to induce protein expression and incubated at 14 °C for 18 h. Cells were harvested by centrifugation at 13,000× g for 20 min at 4 °C and stored at −80 °C for 12 h. Frozen cell pellets were resuspended in 20 mL ice-cold Talon binding buffer (50 mM sodium phosphate pH 7.4 and 300 mM NaCl) containing an EDTA-free protease inhibitor cocktail, and disrupted with a UP50H sonicator (Hielscher Ultrasonic, Teltow, Germany) using 10 cycles (40 s on/60 s off) at 80% pulsar power on the ice. The lysate was centrifuged at 9,400× g at 4 °C for 20 min and the supernatant was applied onto a 5 mL His-Trap TALON crude column (GE Healthcare Bioscience, Piscataway, NJ, USA) pre-equilibrated with deionized water and Talon binding buffer. After sample application, the column was washed with Talon wash buffer (50 mM sodium phosphate pH 7.4, 300 mM NaCl, and 5 mM imidazole) until they reached a steady baseline. Proteins were eluted using Talon elution buffer (50 mM sodium phosphate pH 7.4, 300 mM NaCl, and 150 mM imidazole). The eluted proteins were concentrated using an Amicon Ultra-15 centrifugal filter unit (Millipore, Billerica, MA, USA) at 4,830× g at 4 °C for 15 min. WT and mutant DNA gyrases were further purified by gel filtration chromatography (ÄKTA pure, GE Healthcare Bioscience) using a Hi-Load 16/600 Superdex 200 prep grade column (GE Healthcare Bioscience) equilibrated with 20 mM Tris-HCl (pH 8.0) to remove imidazole. The eluted peaks of the samples (280 nm) were assayed using supercoiling assays and analyzed using SDS-PAGE. Protein concentrations were determined using a Qubit assay kit (Thermo Fisher Scientific, Waltham, MA, USA).

### DNA gyrase activities and inhibition by FQs

DNA supercoiling activity was determined using a combination of purified recombinant *Mul* GyrA and GyrB subunits as previously described (38-40). The reaction mixture (30 µL) consisted of supercoiling assay buffer (35 mM Tris-HCl pH 7.5, 24 mM KCl, 4 mM MgCl_2_, 2 mM DTT, 1.8 mM spermidine, 6 mM ATP, 0.1 mg/mL BSA, 6.5% w/v glycerol) and relaxed pBR322 DNA (0.3 µg) as the substrate. Assays were performed for 1 h at 30 °C and stopped by the addition of 30 µL chloroform/iso-amyl alcohol (24/1) and 3 µL 10× stop and loading solution (40% w/v sucrose, 100 mM Tris-HCl pH 7.5, 1 mM EDTA, 0.5 µg/mL bromophenol blue). The product of the reaction was separated by electrophoresis using a 1% agarose gel in 0.5× Tris-borate-EDTA (pH 8.3) buffer for 90 min at 30 mA, followed by staining the agarose gel with ethidium bromide (0.7 µg/mL). Supercoiling activity was quantified by measuring the band brightness of supercoiled pBR322 DNA using Image J 1.52a (http://rsbweb.nih.gov/ij). A concentration-dependent supercoiling assay using 1 to 24 μM of GyrA or GyrB subunit was performed to determine the optimal concentration of each DNA gyrase subunit. The optimal temperature of WT *Mul* DNA gyrase supercoiling activity was measured at 20, 25, 30, 37, 40, and 50 °C using the same concentrations as stated above and the optimal DNA gyrase subunit concentrations. Inhibition of *Mul* DNA gyrase supercoiling activity by FQs followed previous methods (38-40) with minor modifications. Briefly, reaction mixtures containing optimal DNA gyrase subunits and increasing FQ concentrations (0–320 μg/mL) were assayed as described above. The inhibitory effects of FQs on DNA gyrase activity were assessed by determining the drug concentration required to inhibit the supercoiling activity by 50% (IC_50_) using R studio free software version 1.4.1717 (https://www.rstudio.com). All assays were carried out at least three times and processed on the same day under identical conditions. To facilitate direct comparison, all incubations with WT and mutant DNA gyrase were carried out and processed in parallel on the same day under identical conditions, and assays were done at least three times, with reproducible results. Furthermore, to determine the more detailed functional role of *Mul* DNA gyrases, we performed FQs mediated DNA cleavage assays followed previous methods (32, 33). Supercoiled pBR322 DNA (0.1 μg) was used as the substrate for DNA cleavage assays, and linearized pBR322 DNA by *Hind*III digestion was used as a marker for cleaved DNA. The quinolone concentrations required to induce 25% of the maximum DNA cleavage (CC_25_) were determined for CIP, MOX, and LVX.

### Predicted binding mode between DNA gyrase and FQs by molecular docking

Molecular docking studies and visualization were conducted in the Molecular Operating Environment (MOE 2020.09, Chemical Computing Group ULC, Montreal, Quebec, Canada; https://www.chemcomp.com/index.htm). The docking model was designed using Swiss-Model (https://swissmodel.expasy.org). FQ coordinates were sketched using ChemBioDraw software (PerkinElmer, Waltham, MA, USA), and the amino acid substitutions were generated using the MOE-Protein builder module. Double-stranded oligonucleotides were adopted from the Research Collaboratory for Structural Bioinformatics Protein Data Bank (PDB ID: 6RKS) (37) because our docking model was not included in the double-strand oligonucleotide. Modified docking models were prepared using a flexible docking method with the scores expressed as a sum of five potentials: accessible surface area, Coulomb potential, hydrogen bonds, anisotropy, and van der Waals interactions and refined by MOE through energy minimization. The DNA gyrase and FQ binding energies were estimated using the Amber10: EHT force field and the implicit solvation model of the reaction field was selected. The best binding models were selected for the lowest free energies and optimized RMSD refinement. The distance between amino acid residues on GyrA and the side chain of FQs was calculated using WinCoot-0.9.4.1 (https://bernhardcl.github.io/coot/), and molecular graphics were generated using PyMOL v1.8 (https://pymol.org/2/).

## ACKNOWLEDGEMENTS

We thank Dr. Yuji Miyamoto (Department of Mycobacteriology, Leprosy Research Center, National Institute of Infectious Diseases, Japan) for providing *Mul* cDNA used in this study.

This study was supported partially by funds No. 21fk0108139j2302, JP18fk0108064, JP18fm0108008, JP18fk0108042, JP18jk0210005, and JP18jm0510001 from the Japan Agency for Medical Research and Development (AMED), partially by a Grant-in-Aid for Young Scientists (B) from the Japan Society for Promotion of Science (No. 17K15695), a grant from the Ministry of Education, Culture, Sports, Science, and Technology (MEXT) of Japan. The funders had no role in study design, data collection and analysis, decision to publish, or preparation of the manuscript.

